# *MapMyCells:* High-performance mapping of unlabeled cell-by-gene data to reference brain taxonomies

**DOI:** 10.64898/2026.03.06.710160

**Authors:** Scott F. Daniel, Changkyu Lee, Tyler Mollenkopf, Matthew Lee, Joel Arbuckle, Elysha Fiabane, Mariano I. Gabitto, Nelson Johansen, Inkar Kapen, Andrew W. Kraft, Jane Lai, Su Ying Li, Ryan McGinty, Jeremy A Miller, Skyler Welch-Moosman, Sven Otto, Lane Sawyer, Noah Shepard, Carol L. Thompson, Andreas Tjärnberg, Jack Waters, Xingjian Zhen, Evan Macosko, Ed S. Lein, Lydia Ng, Hongkui Zeng, Shoaib Mufti, Zizhen Yao, Michael J. Hawrylycz

## Abstract

Single-cell mapping methods convert raw, heterogeneous single-cell datasets into interpretable and comparable representations of biological identity. As reference cell-type taxonomies mature, mapping new datasets to shared references has become a central strategy for enabling cross-study integration, reproducible annotation, and cumulative biological knowledge. Here we present *MapMyCells*, an open-source framework designed to align diverse single-cell omics datasets to hierarchical reference taxonomies with minimal preprocessing. MapMyCells provides out-of-the-box support for an expanding set of high-quality brain cell-type references generated by the Allen Institute for Brain Science, the BRAIN Initiative, and the Seattle Alzheimer’s Disease Brain Cell Atlas, including whole-brain mouse and human atlases, aging and Alzheimer’s disease cohorts, and a cross-species consensus taxonomy initially focused on the basal ganglia. MapMyCells enables efficient mapping of hundreds of thousands of cells on standard workstations without specialized hardware, providing a deterministic, scalable, and modality-agnostic approach that is robust across species and molecular assays. The framework produces interpretable confidence metrics and quantitative summaries of mapping performance, allowing users to evaluate assignment precision and accuracy. We demonstrate the mapping of unlabeled transcriptomic, epigenomic, and spatial datasets to reference taxonomies and describe a general workflow for preparing arbitrary hierarchical taxonomies for reference-based mapping. As the ecosystem of single-cell reference atlases expands, MapMyCells offers a practical and reproducible solution for community-scale cell-type annotation and cross-dataset integration, supporting the development of unified and extensible brain cell atlases.

## 1. Cell type mapping in single cell genomics

Single-cell transcriptomics has transformed neuroscience by generating large-scale, high-dimensional datasets that capture molecular diversity across tissues, developmental stages, and species. The central challenge is no longer data acquisition but the establishment of coherent reference frameworks that enable individual experiments to contribute to cumulative and reusable biological knowledge. Recent advances in single-cell genomics and computational analysis have enabled the construction of data-driven cell-type taxonomies grounded in molecular signatures, providing a principled and quantitative complement to classifications defined by classical anatomical or physiological criteria^1–4^. These taxonomies formalize cell identity within a testable framework, enabling consistent comparison across studies, platforms, and species while supporting both basic discovery and translational applications^5,6^.

A major conceptual shift accompanying this progress is the emergence of reference-based analysis. Rather than analyzing each dataset independently through unsupervised clustering, increasingly mature reference atlases now allow new data to be interpreted in the context of accumulated prior knowledge. This transition parallels the impact of the first human genome references, which transformed genomics from isolated studies into a cumulative and comparative science^6^. In reference-based workflows, annotated atlases define shared latent representations into which query datasets can be projected, enabling consistent cell-type identification across disease states, molecular modalities, perturbations, and species^57^ .

At the core of this paradigm is *cell-type mapping*, whereby reference taxonomies serve as templates for aligning, annotating, and integrating newly generated datasets. Anchoring data to a shared reference improves interpretability, promotes cross-study harmonization, and provides a scalable solution as dataset size and complexity continue to grow. Beyond routine annotation, mapping establishes a quantitative framework for measuring biological concordance and divergence across experimental conditions, developmental trajectories, and evolutionary contexts. Importantly, reference mapping is not a static process: as new datasets are incorporated, taxonomies themselves can be iteratively refined, allowing cell-type definitions to evolve while preserving continuity with prior knowledge. This iterative refinement is essential for building stable yet adaptable frameworks capable of supporting large-scale collaborative initiatives, comparative biology, and downstream applications such as disease modeling, target discovery, and precision therapeutics.

Operationally, reference-mapping workflows typically involve three conceptual stages: 1) learning a shared representation that preserves biological structure while mitigating technical variation; 2) associating reference cells with curated or ontology-based annotations; and 3) transferring discrete labels or continuous features to query datasets through learned alignment functions^6,7^. Automated classification and alignment algorithms thus serve as engines for translating prior biological knowledge into reproducible annotations. When properly applied, these approaches reduce dependence on repeated reclustering and manual curation, enabling scalable analysis and more reproducible integration across experiments, modalities, and species.

The scale and heterogeneity of contemporary datasets, however, make manual cell-type annotation increasingly impractical and inherently subjective. The integration of diverse experimental designs, sequencing technologies, and molecular modalities therefore requires robust computational strategies. As a result, a broad ecosystem of mapping approaches has emerged (**Fig. 1A**), spanning methods based on marker genes and linear embeddings^8–11^, supervised classifiers trained on reference atlases^12–14^, probabilistic generative models that explicitly model cell-state uncertainty and mixture structure^15,16,17^, and latent-space alignment frameworks using matrix factorization, variational inference, and deep learning^18–20^. Despite rapid progress, critical challenges remain, including reliable identification of novel or rare cell types, preservation of genuine biological variation while suppressing technical noise, and the development of interoperable ontological standards that allow annotations to remain comparable across atlases. Addressing these challenges will be central to establishing unified, extensible brain cell atlases capable of supporting both mechanistic neuroscience and translational research.

**Figure 1.**
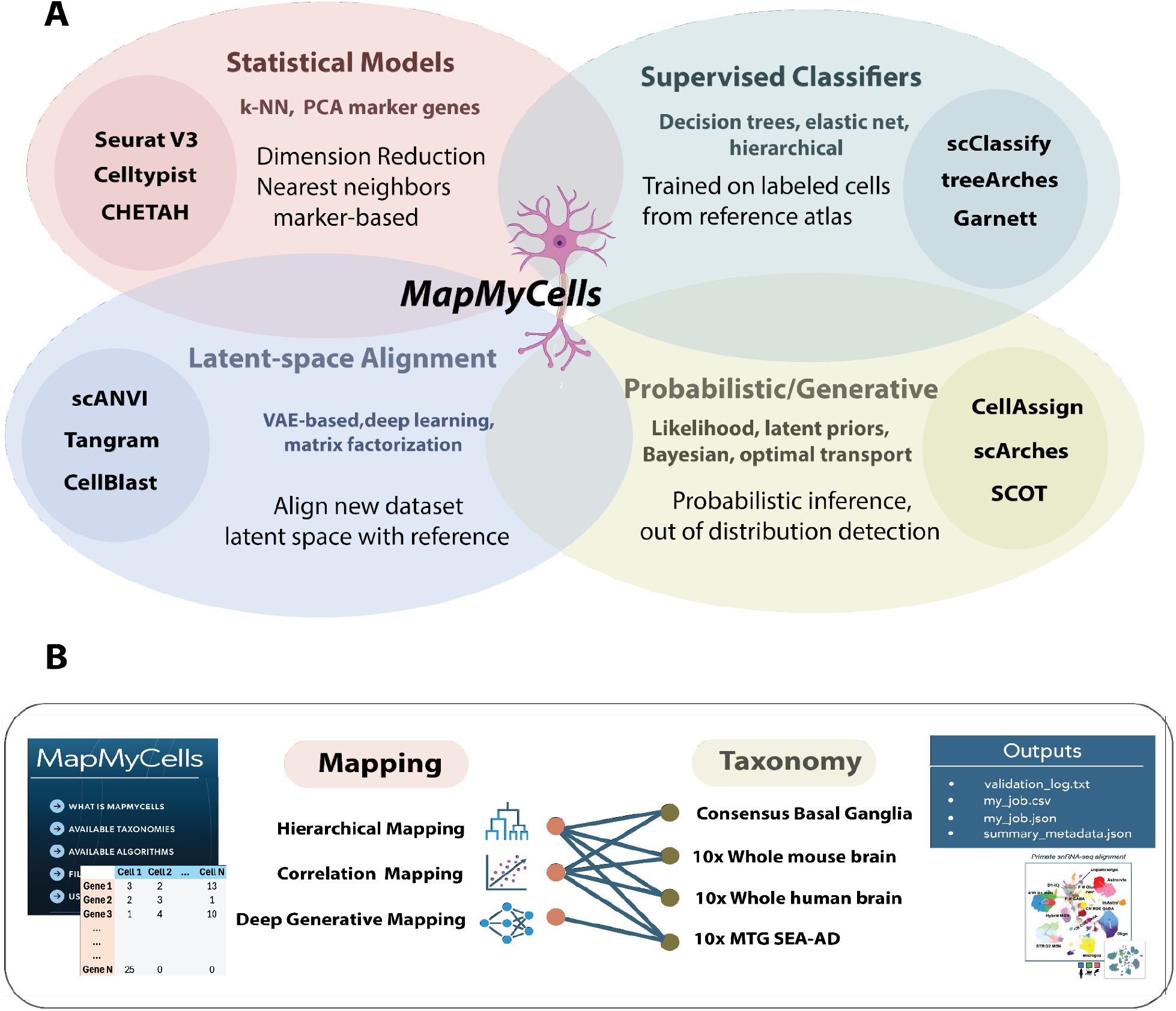
Single cell mapping approaches for Omics data. A) A wide variety of methods have been devised for mapping single cell RNA-seq and epigenetic data. While there is no unique characterization of the methods, major techniques include methods from classical statistical models leveraging nearest neighbor mapping or canonical correlation (Seurat^21^, Cell Typist^9^, CHETAH^11^), supervised classifiers trained on labelled cells using reference atlases (scClassify^12^, treeArches^14^, Garnett^13^), probabilistic/generative methods using likelihood or Bayesian priors and advanced optimization methods such as optimal transport (CellAssign^16^, scArches^22^, SCOT^15^). Latent-space alignment methods use latent space reduction, variational autoencoders and methods from deep learning (scANVI^19^, Tangram^18^, CellBlast^20^). MapMyCells is a suite of algorithms using techniques from basic statistics, hierarchical reference atlas, and variational autoencoders. The MapMyCells uses techniques from statistical models, supervised classifiers, and latent space alignment. B) MapMyCells references include a set of multi-species taxonomies for mouse and human, a multimodal multispecies basal ganglia taxonomy generated using human, macaque, and marmoset, and a modified middle temporal gyrus (MTG) human cortical taxonomy which includes Alzheimer’s disease vulnerable types. Currently, three mapping methods are available include correlation based, hierarchical, and a variational autoencoder model^3^. (See **Section 2** and **Methods**)

The Allen Institute for *Brain Science Brain Knowledge Platform (BKP)*^*23*^ is an ecosystem of interoperable tools designed to describe, relate, and operationalize cell-type taxonomies derived across species, brain regions, and experimental modalities. Within this ecosystem, *MapMyCells* provides a unified framework for mapping unlabeled single-cell datasets onto predefined reference taxonomies^24^. The framework integrates label-transfer strategies that combine classical statistical approaches, supervised classification, and latent-space alignment, with less emphasis on probabilistic/generative models (**Fig.1A**) with an emphasis on usability, computational efficiency, and broad accessibility to community reference resources.

A defining strength of MapMyCells is its scalability. Leveraging cloud-enabled infrastructure within the Brain Knowledge Platform and reference datasets comprising millions of cells, MapMyCells supports mapping workflows at whole-brain and large-cohort scale, including datasets containing up to hundreds of millions of cell–gene measurements. By enabling direct comparison of user-generated data with high-quality, high-resolution reference taxonomies, MapMyCells accelerates biological interpretation, supports consistent cross-study integration, and facilitates the incorporation of community-generated datasets into shared taxonomic frameworks.

MapMyCells is available both as a web-based application^24^, providing one-click access to Allen Institute– supported reference taxonomies, and as an open-source Python library^25^ that enables flexible ingestion of user data and mapping to custom-defined hierarchies. The application is built on cloud infrastructure to ensure reliability and scalability, while the lightweight library supports deployment on high-performance computing clusters, cloud environments, or standard local workstations. Importantly, memory usage scales primarily with taxonomy complexity rather than raw dataset size, making the framework well suited for references derived from multi-million-cell transcriptomic studies. In the following sections, we describe and evaluate MapMyCells. **Section 2** outlines currently available reference taxonomies and associated mapping strategies, including marker-selection procedures. **Section 3** demonstrates application of MapMyCells to a whole-mouse-brain reference taxonomy and evaluates mapping performance across multiple modality datasets and taxonomies with respect to approximate ground truth. Finally, **Section 4** benchmarks MapMyCells against alternative methods in a representative end-to-end workflow. Together, these analyses show that MapMyCells provides accurate, scalable label transfer without requiring specialized hardware or large-memory systems, lowering practical barriers to adoption across the neuroscience community.

## 2. *MapMyCells* Taxonomy and Mapping

After more than a century of neuroanatomical and physiological investigation, it is now firmly established that neurons and glial cells, like cells in other tissues, comprise diverse and reproducible cell types ^26^. Whereas early classifications have relied primarily on morphology or electrophysiological properties^26–28^, contemporary brain cell type taxonomies instead seek to define biologically meaningful and quantitatively reproducible units of cellular diversity using high-dimensional molecular measurements augmented by other data modalities. Recent taxonomies have been constructed using single-cell transcriptomics^1,29^, epigenomics ^1,29,30^, spatial transcriptomics ^31^, and integrative analyses incorporating morphology and electrophysiology^3233^. Although cell identity often exhibits continuous variation and context dependence, these representations are commonly organized as hierarchical taxonomies. Such hierarchies provide a practical and computationally convenient framework for naming, comparison, and cross-study harmonization^32,34^; however, the underlying biological structure need not be strictly tree-like and may instead reflect manifold structure, gradients, or partially ordered relationships^35^.

### 2.1 MapMyCells Taxonomies

To address the intrinsic complexity of brain cell identity, encompassing species specific and regional variation, discrete classes, continuous gradients, developmental trajectories, and state-dependent variation, multiple complementary taxonomies and alignment methods are required. The MapMyCells pipeline (**Fig. 1B**) currently supports four reference taxonomies spanning species and brain regions. Each taxonomy undergoes systematic validation through marker gene enrichment analysis, cross-dataset reproducibility/interpretability and structural robustness across levels of the organizational framework.

The four reference taxonomies presently supported by MapMyCells include whole mouse brain (WMB, CCN20230722 ^1^), Whole Human Brain (WHB, CCN20240330^2^), Cross-species consensus basal ganglia (CBS, CCN20250428^36^), and Human medial temporal gyrus, Alzheimer’s disease tissue (MTG, CCN20230505^3^) (**Box 1**). Each of the four taxonomies are ingested into the web application through certain preprocessing steps that take the single cell data defining the taxonomies and reduce it into artifacts required by MapMyCells. The Python library^25^ provides tools to perform this preprocessing so that interested users running the Python code locally can ingest and map to any arbitrary cell type taxonomy. We discuss this process in **Section 3** below and in the **Methods** section.

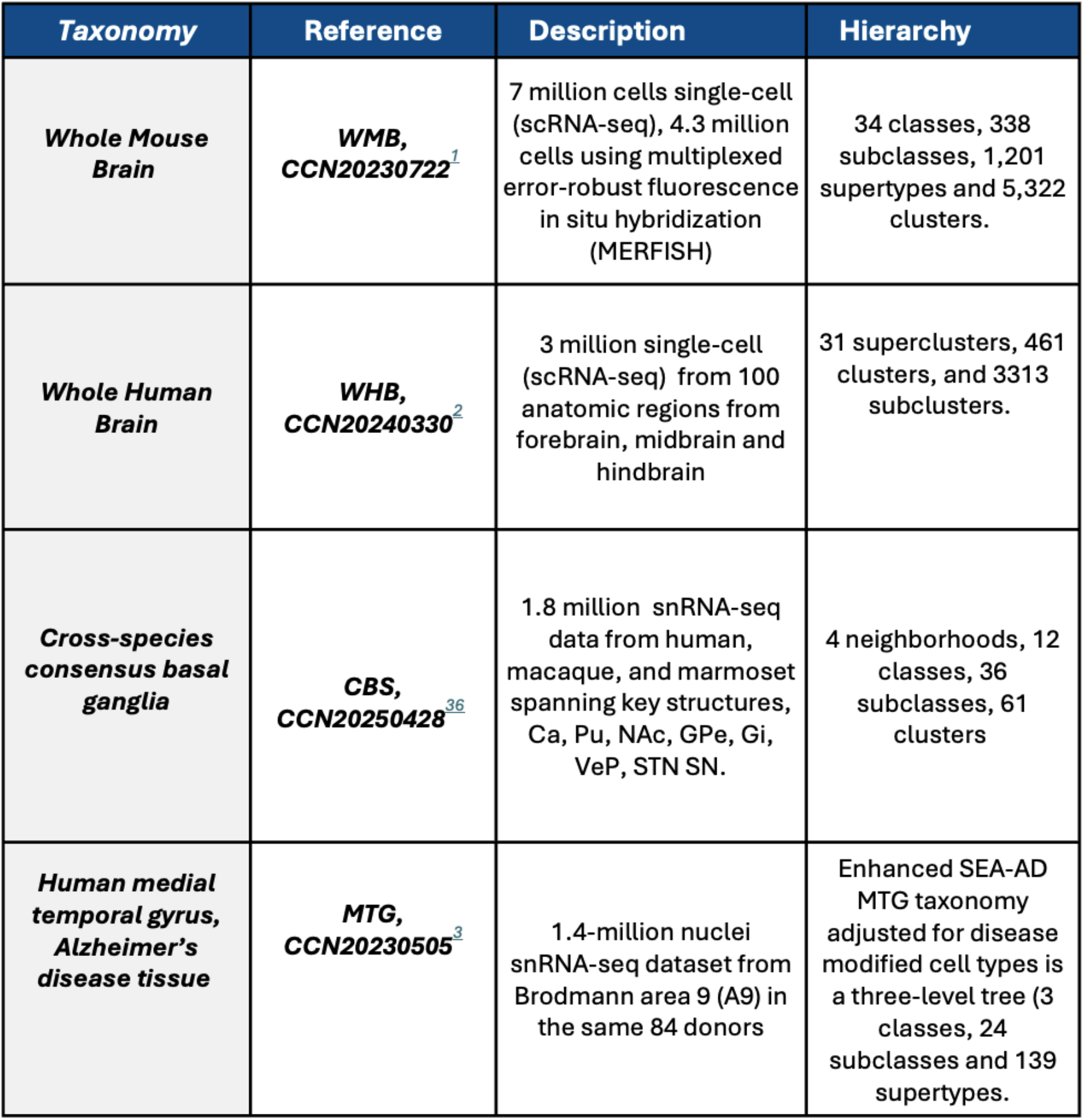
Box 1. MapMyCells Taxonomies. Four taxonomies presently supported by MapMyCells: Access/Reference provides reference for online use; Size/Description is scale and modality; Hierarchical structure gives nomenclature description. Basal ganglia abbreviations Caudate (Ca), putamen (Pu), nucleus accumbens (NAc), external and internal globus pallidus (GPe, GPi), ventral pallidus (VeP), subthalamic nucleus (STN), and substantia nigra (SN).

### 2.2 Mapping algorithms

Brain cell mapping techniques now represent a wide range of scalable mapping algorithms that enable robust integration and comparison of large, heterogeneous single-cell and spatial transcriptomic datasets across studies, modalities, and species^6^. MapMyCells was developed as part of BRAIN Initiative Cell Atlas Network (BICAN)^37^ and currently offers a choice of correlation based, hierarchical mapping, or a deep hierarchical algorithm based on conditional variational autoencoders. The more basic mapping algorithms of correlation and hierarchical mappings are designed for high performance use. Because these methods rely solely on classical statistical algorithms, they perform robustly without specialized hardware or extensive memory.

The most basic mapping algorithm is *correlation mapping*, a one step nearest cluster centroid mapping based on correlation. For a given taxonomy having a cell by gene count matrix and assigned clusters, each reference cluster is summarized by cluster mean and its marker genes **(Methods**). The cluster assignment is made for query data by selecting the cluster of the minimal distance (cosine distance) to the query data using pre-calculated marker genes with correlation as distance metric. Correlation mapping is a reasonable option when data is generated by the same sequencing platform as the reference data being mapped against. It is robust for most single-cell RNA-Seq (10X v3, 10X v2, SMART-Seq) and MERFISH datasets, and is faster and simpler than other methods.

The MapMyCells *hierarchical* algorithm was originally developed to map spatial transcriptomics data onto the hierarchical cell type taxonomy derived from single cell transcriptomics data^1^ and is suitable for both cross-species and cross-platform use. The method can be used to map to any taxonomy provided the taxonomy is strictly hierarchical. All taxonomies described in **Section 2** are hierarchical where the underlying graphical structure is a tree without loops or multiple edges. Internally the algorithm uses two sets of data: 1) A file specifying the average gene expression profile of each cell type in the taxonomy, and 2) A file specifying marker genes at each parent cell type (i.e. each cell type with more than one child cell type) in the taxonomy. Internal to each taxonomy, marker genes associated with a given cell type are used to discriminate between cell types that are children of the type. MapMyCells is agnostic to how these marker genes are selected, and the MapMyCells Python library provides functions to implement an algorithm for marker gene selection for a new taxonomy.

The MapMyCells hierarchical algorithm is described in **Methods and Suppl. Fig. 1**. Briefly, each cell is assigned to the root as the current node. In descending the tree toward leaf types, at a given node one of the child types will be selected. To decide, the user input cell expression vector is correlated over all leaf types (lowest level of taxonomy) descended from the current node using the differentiating markers at the current node. A bootstrap sampling of the markers is used to assign a likelihood score to the best correlated child of the current node. Bootstrapping is repeated 100 times. This child type is then determined to be the best fit cell type from the plurality of iterations and becomes the new current node.

Two metrics of mapping strength and uniqueness are inherent in the hierarchical algorithm (**Methods**): 1) the average correlation coefficient of a cell to its assigned cell type, and 2) the fraction of bootstrap iterations that assigned the cell to that type. While both correlate with the accuracy of the cell type mapping, the *bootstrap probability* (the fraction of iterations that chose the assigned type), provides a better indication of whether an assignment is trustworthy (**Supp. Figure 1**). The average correlation coefficient is a useful metric for identifying out-of-distribution cells. One deficiency of the MapMyCells hierarchical algorithm is that every cell must be assigned to a cell type in the taxonomy. If a cell is assigned to a cell type, but is better represented by a type outside of the chosen taxonomy, this will be exposed by lower values in the strength and uniqueness metrics.

The third mapping algorithm is *Deep generative mapping* and is an extension of conditional variational autoencoders ^3^ methods. It is described here briefly and will be described in detail in a future publication. Deep Generative Mapping is a generative model for mapping snRNA-seq data sets and assigning cell type identity. Cells mapped with this method are not only assigned an associated cell type, but also associated confidence on mapping. The algorithm can be used on novel snRNA-seq data set generated in human tissue to assign cellular identity, and associated confidence intervals on mapping, based on Seattle Alzheimer’s Disease Brain Cell Atlas (SEA-AD). The model uses a variational autoencoder neural network, which first embeds the original input data into latent space and then trains a multilayer perceptron (MLP) model to further classify the labels. In the last layer of the MLP classifier, there is a SoftMax layer that normalizes across all labels into probabilities. The deep generative mapping algorithm has been benchmarked on 10x Human MTG SEA-AD (CCN20230505) data with 80% training and 20% as evaluation: supertype level (for testing set): mean F1=0.86, median F1=0.90, accuracy=0.91. Subclass-level: mean F1=0.99, median F1 = 0.996, accuracy=0.995.

### 2.3 Use Case: Mapping to Whole mouse brain taxonomy

We illustrate the use of MapMyCells by mapping single cell and spatial transcriptomics data sets to a whole mouse brain taxonomy^1^. We developed an online site^38^ that includes an R workbook illustrating this common use case for the MapMyCells facility. The query data included for this vignette are single cell and spatial transcriptomic data from another major whole brain atlas developed through BICAN^39^. **Figure 2** presents a typical workflow. After downloading the data sets, loading relevant libraries, and performing some basic QC, we transfer labels from Whole Mouse Brain taxonomy *(WMB, CCN20230722)* onto the user input droplet-based sequencing cells using MapMyCells, and then visualize the results in a UMAP. **Fig. 2A** shows the input requirements and file size limits for data input either as .csv or .h5ad file types. H5ad files are produced by AnnData^40^, a widely used tool for creating, manipulating, and saving large data matrices, such as for expression data.

**Figure 2.**
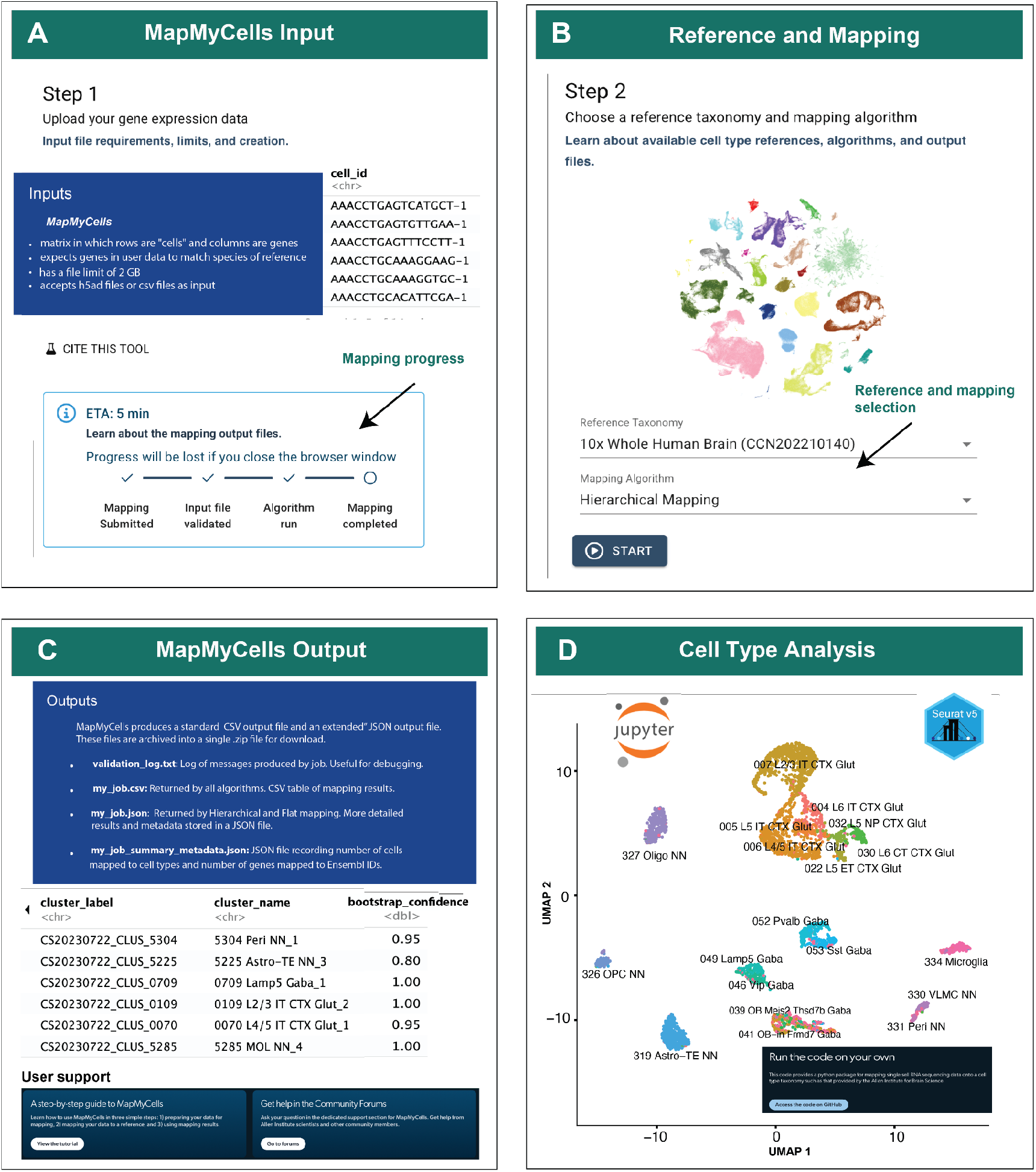
Mapping to Whole Mouse Brain Taxonomy. A) MapMyCells interface guides users through the selection and mapping process. Step 1 determines input data formats and size requirements. Once mapping begins a progress bar tracks the process. B) The user selects taxonomy and mapping methods to start mapping. C) Output is a set of files bundled into a single .zip file for download. In addition to the mapping results, log files summarize the mapping process for debugging. File my_job.csv has a line for each cell and for each level of mapped resolution the type name and its bootstrap confidence (**Methods**.) Extensive online documentation provides help, examples, and user support. D) Following download a variety of tools can be used to analyze results. Shown is a UMAP visualization in Seurat of mapped mouse mapped cells to subclass level. MapMyCells documentation provides access to libraries and documentation for offline use.

MapMyCells online has a file limit of 2 GB. Code for compressing data and for splitting data set into multiple input files are provided^38^ or the users can run MapMyCells offline^25^ with no size restrictions.

In **Fig. 2B** we select the reference taxonomy and mapping algorithm to use, in this case *hierarchical mapping*. Once mapping is started a progress bar (left) will inform about status. When mapping is complete there is an option to download the output as a compressed file with three files: “validation_log.txt”, “[#].json”, and “[#].csv”. The key output of the mapping is [#].csv which contains the mapping results while the validation .txt and .json files provide information about the mapping status (**Fig. 2C**). The file .csv output contains cluster ids, labels at each level of mapping hierarchy (e.g. subclass, supertype, cluster) and corresponding bootstrap probabilties (i.e. fraction of bootstrapping runs that the cell mapped to each class). Additional user help is provided online. To visualize the results in R, convert the data into a Seurat^41^ object with (modified) mapping results as metadata and run the standard pipeline for creating a UMAP in Seurat, and color-code each cell. Here the data (**Fig. 2D**) has not been clustered but rather the subclass assignments are assigned to the data in the UMAP. The subclasses shown are largely segregated from one another without the need to apply clustering, suggesting this mapping works well at this resolution.

## 3. Measuring MapMyCells performance

We present several studies of MapMyCells’ performance, focusing specifically on the hierarchical mapping algorithm. In Section 3.1, we perform test-train holdout evaluations on the datasets used to derive the original taxonomies which MapMyCells supports and show that MapMyCells is able to reproduce the ground truth annotations of these datasets with a high degree of accuracy. In Section 3.2, we evaluate MapMyCells’ performance on an external ATAC seq dataset^42,43^ by mapping it to the Whole Mouse Brain taxonomy^1^. Inspection of the resulting cell type distributions shows that MapMyCells has produced credible cell type mappings. In Section 3.3, we evaluate performance on data with a spatial component by mapping an independently generated spatial transcriptomics dataset^44^. Finally, in Section 3.4, we summarize a novel use case in which MapMyCells is used to assign cells from a recent BICAN basal ganglia dataset to anatomical zones based only on their transcriptomic signatures.

### 3.1 Mapping data used to derive taxonomies

The most straightforward evaluation of a label transfer algorithm is to take the data used derive a taxonomy, split the data into a large training and a smaller test set, train the algorithm using the training set (in the case of MapMyCells, “training” means finding marker genes and deriving the mean gene expression profiles for cell types), and then map the test set back onto the taxonomy. Since the test set is part of the data used to derive the taxonomy, there is in principle ground truth to which we can compare the results. We perform this test-train holdout evaluation on three BICAN taxonomies: the Yao et al. 2023 Whole Mouse Brain taxonomy ^1^, the Siletti et al. 2023 Whole Human Brain taxonomy ^2^, and the Whole Mouse Brain taxonomy from Langlieb et al. 2023^45^. This last taxonomy is presently not one of the taxonomies incorporated as a reference taxonomy in MapMyCells. For that, we downloaded the publicly available data used to derive the Langlieb et al. taxonomy^46^, preprocessed it, and mapped the test set using the open-source Python library supporting MapMyCells^25^.

**Figure 3A** shows the performance of MapMyCells at each level (e.g. class, supertype) of these test taxonomies. The horizontal axis is the threshold bootstrap probability metric, representing whether a cell’s mapping is considered valid. At higher levels of the taxonomy (e.g. class, subclass) MapMyCells performs very well, achieving macro averaged F1 values (upper) between 0.99 and 0.90, and with high fractions (lower) of correctly labelled cells. However, this evaluation is less realistic as it assumes that users are using an unlabeled dataset with the exact same signal to noise properties and gene panel as the training set (i.e. all marker genes are present). Although simulating different signal-to-noise properties is difficult, simulating different gene panels is easy, and especially useful when considering performance on spatial transcriptomics data, which tends to have much smaller gene panels than single cell or single nucleus RNA sequencing data.

**Figure 3.**
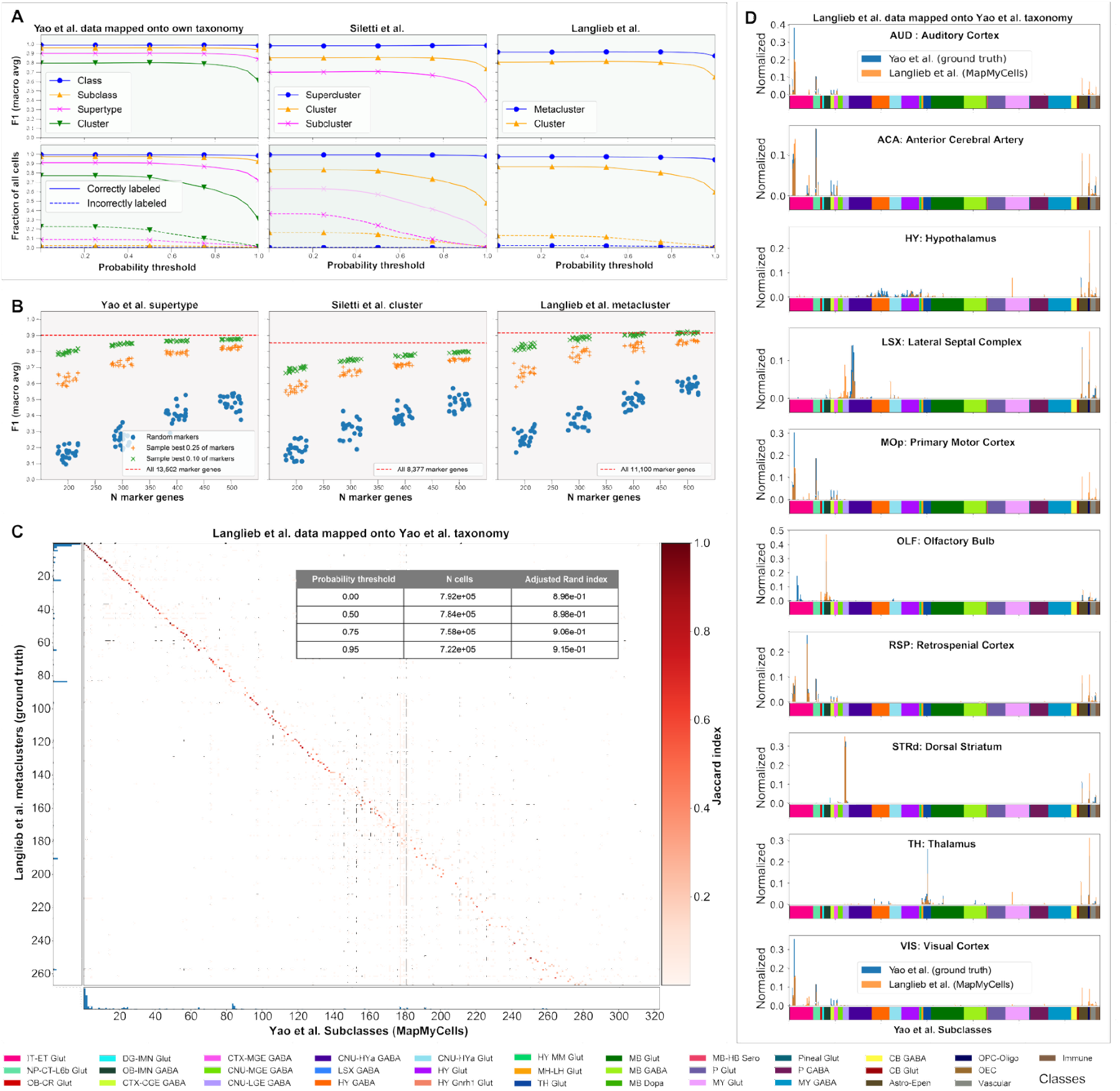
Mapping within modality. A) The result of mapping data used to derive each taxonomy back onto that same taxonomy. Performance metrics as a function of quality cuts in bootstrapping probability for select levels of supported taxonomies. Averaged F1 values (upper) between 0.90 and 0.99 for the highest level of the taxonomy, and high fractions of correctly labelled cells (lower). B) Degradation of performance as the gene panels are artificially downsampled to contain fewer and fewer marker genes. Blue circles are randomly selected gene panels. Orange ‘+’ are randomly selected from the most discriminating 25% of marker genes. Green ‘x’ are gene panels sampled from the most discriminating 10% of marker genes. C) Jaccard index of intersecting Broad metaclusters and Allen subclasses found by MapMyCells. A strong diagonal component confirms an underlying similar biological structure of the data. The inset table shows the adjusted Rand index quantifying the overall equivalence of the two annotations. D) Similarity in distribution of cell type subclasses from ten Allen anatomic regions (AUD, ACA, HY, LSX, MOp, OLF, RSP, STRd, TH, VIS) with the cell types mapped from Broad data taken from those regions. Cell type class color code shown below.

**Figure 3B** performs the same test-train holdout evaluation, but artificially downsamples the test data so that it does not contain every marker gene that MapMyCells expects. Downsampling the gene panel in a purely random way (blue circles) significantly degrades mapping performance. However, if we assume that gene panels are used which at least contain a fraction of the best markers we see that F1 performance is essentially unaffected. The orange ‘+’ and green ‘x’ symbols in **Fig. 3B** presents the downsampled gene panel evaluation by intentionally selecting the downsampled gene panel from only the most powerful markers, i.e. those that differentiate the largest number of cell types. Here,MapMyCells is able to map datasets with as few as 400 marker genes nearly as accurately as datasets containing the full complement of several thousand markers.

A more challenging experiment is to map data used to define one taxonomy onto another taxonomy, and compare the annotations of each cell in each taxonomy for credible biological correspondence. **Figures 3C, 3D** show the result of mapping two whole mouse brain taxonomies where the^1,45^ ∼5,000 clusters of the Langlieb et al.^39^ taxonomy are mapped onto the Allen ^1^ taxonomy of comparable size. The mapping process (training on the taxonomy and then mapping the test set) took 5.5 hours and 70 GB of memory on a 16 core workstation. **Fig. 3C** shows the Jaccard index intersecting Broad metaclusters and Allen subclasses found by MapMyCells. A strong diagonal component confirms an underlying similar biological structure of the data. The inset table shows the adjusted Rand index quantifying the overall equivalence of the two annotations which improves as more stringent cuts on the quality of cell mappings are accepted as valid. To more closely examine the mapping, in **Fig. 3D** we subdivide the dissections into ten major cortical and subcortical regions (AUD, ACA, HY, LSX, MOp, OLF, RSP, STRd, TH, VIS) from which samples from Broad and Allen data were independently taken. We then compare the distribution of subclasses found by mapping the Broad data with the original subclasses defining each region in the Allen data. In each region, the plots show a similar distribution of Allen (blue) defining subclasses with mapped Broad (orange) subclasses indicating a common cell type structure identified by each assay.

MapMyCells’ successful performance in analyzing and mapping these two whole brain datasets facilitated developing the first *Consensus Whole Mouse Brain (WMB, CCN202510310)*^*47*^ integrated taxonomy, built upon Allen Institute single-cell RNA sequencing (scRNA-seq) and Broad Institute single-nucleus RNA sequencing (snRNA-seq) data. The Broad Institute taxonomy, constructed from 4.4 million snRNA-seq profiles, defines 16 classes, 223 metaclusters, and 5,030 clusters^45^. Integrating these two large-scale taxonomies into a unified framework represents a consensus view of cell types from independent assays and laboratories across the entire mouse brain.

### 3.2 Mapping across modalities

Modern cell-type taxonomy construction includes epigenomic assays such as ATAC-seq, DNA methylation profiling, and other measurements of chromatin accessibility and regulatory dynamics^43,48^. Direct comparison and mapping between these modalities remain challenging due to inherent modality mismatches, uncertainties in peak-to-gene assignments, and differences in sparsity and signal-to-noise characteristics. Nevertheless, developing approaches to compare and align taxonomies derived from these diverse modalities is essential, as each captures complementary aspects of gene regulation and cellular function and therefore contributes distinct biological insight into cell-type identity^30^.

To evaluate the performance of MapMyCells on cross modality data, we downloaded mouse brain ATAC seq data from a study of gene regulatory elements in the mouse cerebrum^43,49^. This data consists of approximately 800,000 cells collected from the isocortex, olfactory bulb, hippocampus, and cerebral nuclei of adult mice and sequenced via single nucleus ATAC-seq^43,49^. The authors clustered this data to obtain an annotated three level taxonomy consisting of 3 classes, 43 subclasses, and 160 cell types. We evaluate MapMyCells’ performance by mapping this data to the Whole Mouse Brain taxonomy (WMB) and examining both the distribution of cell types per anatomical dissection and the correspondence between the WMB and the ground truth ATAC-seq derived taxonomy. No special preprocessing was done to account for the differences between ATAC-seq and RNA-seq data; we downloaded the data from CZI cellxgene^49^ and mapped it with the MapMyCells Python library^25^.

Figure 4 shows the result of mapping to the WMB taxonomy. **Fig. 4A** shows (similarly to **Fig. 3D**) the distributions of WMB subclasses in several anatomical regions of the Li et al.^43,48^ data. Comparable regions were identified in both parcellations and included pallidum (PAL), paracentral lobule-infralimbic area-orbital cortex (PL-ILA-ORB), lateral septal complex (LSX), and olfactory bulb (OLF). Here the distribution of subclasses mapped (orange) agrees well with the expected subclass distribution of WMB. One exception are cells spuriously assigned to subclasses descended from the *MY Glut* class in each anatomical dissection, which is a class containing poorly defined subclasses comprising low quality cells. (**Suppl. Fig.2**) Applying a quality cut to the mapping result suppresses the presence of these spurious MY Glut cells resulting in the green histograms of **Fig. 4A** and after-cut confusion matrix (**Fig. 4B)** relating the cell type WMB classes found by MapMyCells with the subclasses derived from the epigenetic data. The strong diagonal illustrates that cells of major GABAergic, glutamatergic, and non-neuronal classes are all appropriately assigned.

**Figure 4.**
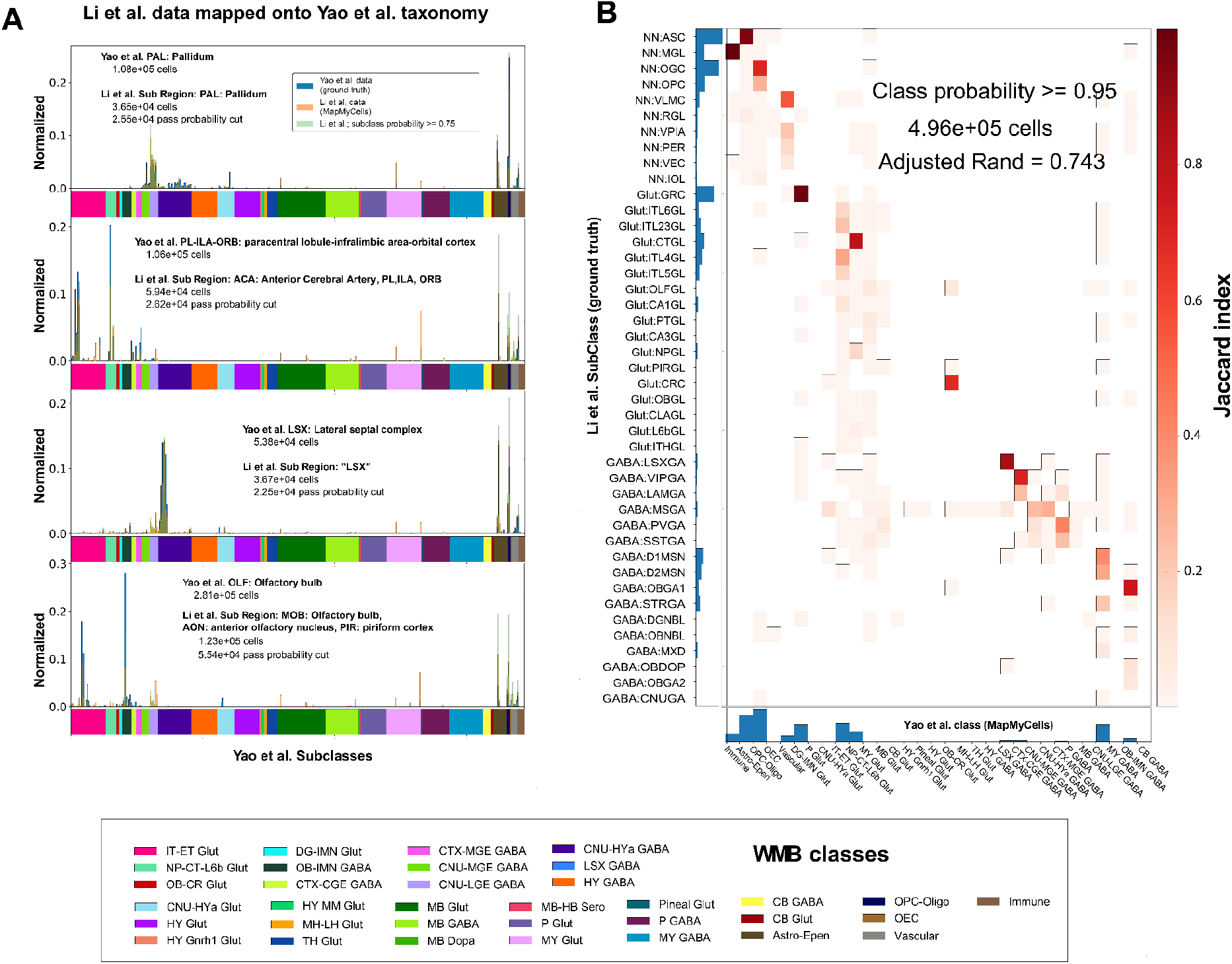
Cross modality mapping. A) Single nucleus ATAC-seq data^43,49^ mapped to the Whole Mouse Brain taxonomy (WMB). Comparing distribution of subclasses found by MapMyCells (orange) to those in the WMB (blue) across different anatomical dissections identified as corresponding in both datasets. PAL: pallidum, PL-ILA-ORB: paracentral lobule-infralimbic area-orbital cortex, LSX: lateral septal complex, OLF: olfactory bulb. B) Histogram and confusion matrices (after quality control cut) relating the cell type WMB classes found by MapMyCells with the subclasses with which Li et al^43,49^. annotated their data. Correspondence improves from adjusted Rand 0.47 to 0.743 when a quality cut is applied to the mapped cells. WMB class annotation colors shown below.

### 3.3 Mapping spatial transcriptomics data

Spatial transcriptomics measures gene expression *in situ*, preserving the anatomical, cellular, and circuit context of the tissue, directly linking molecular identity to spatial organization of cell types^50^ . Automatic label transfer in this context enables matching single cell RNA-seq data to data with a resolved spatial context. To test MapMyCells’ performance, we downloaded publicly available spatial transcriptomics data from a spatial atlas of the mouse central nervous system^44^ which used the StarMAP PLUS spatial transcriptomics platform^51^ to sequence 1.1 million cells in 1,022 genes across 20 anatomical slices covering the entire mouse central nervous system. **Figures 5A,B** show the spatial distribution of cell types mapped by MapMyCells across four representative anatomical slices. The cell-by-gene table was directly mapped without additional preprocessing, and the spatial transcriptomics gene panel was selected independently of the MapMyCells marker gene panel. Assigned cell types qualitatively agree with cell types assigned to these anatomic structures determined by spatial transcriptomic data in WMB^31,50^. Class level annotation of the sagittal section in **Fig. 5A** shows major cell types in midbrain (MB Gaba, MB Dopa), cerebellum (CB GABA, CB Glut), hypothalamus (HY Glut, HY Cnrh1 Glut), striatum (LSX GABA, CNU-LGE GABA), cortex (IT-ET Glut), and olfactory bulb (OB IMN GABA), while a refined subclass level mapping (**Fig. 5B**) shows increasing resolution in all structures including cortex (L2/3 IT CTX Glut, L4/5 IT CTX Glut), hippocampus (CA1-ProS Glut, CA3 Glut). The complex mixture of cell types in lower brain structures is also apparent.

**Figure 5.**
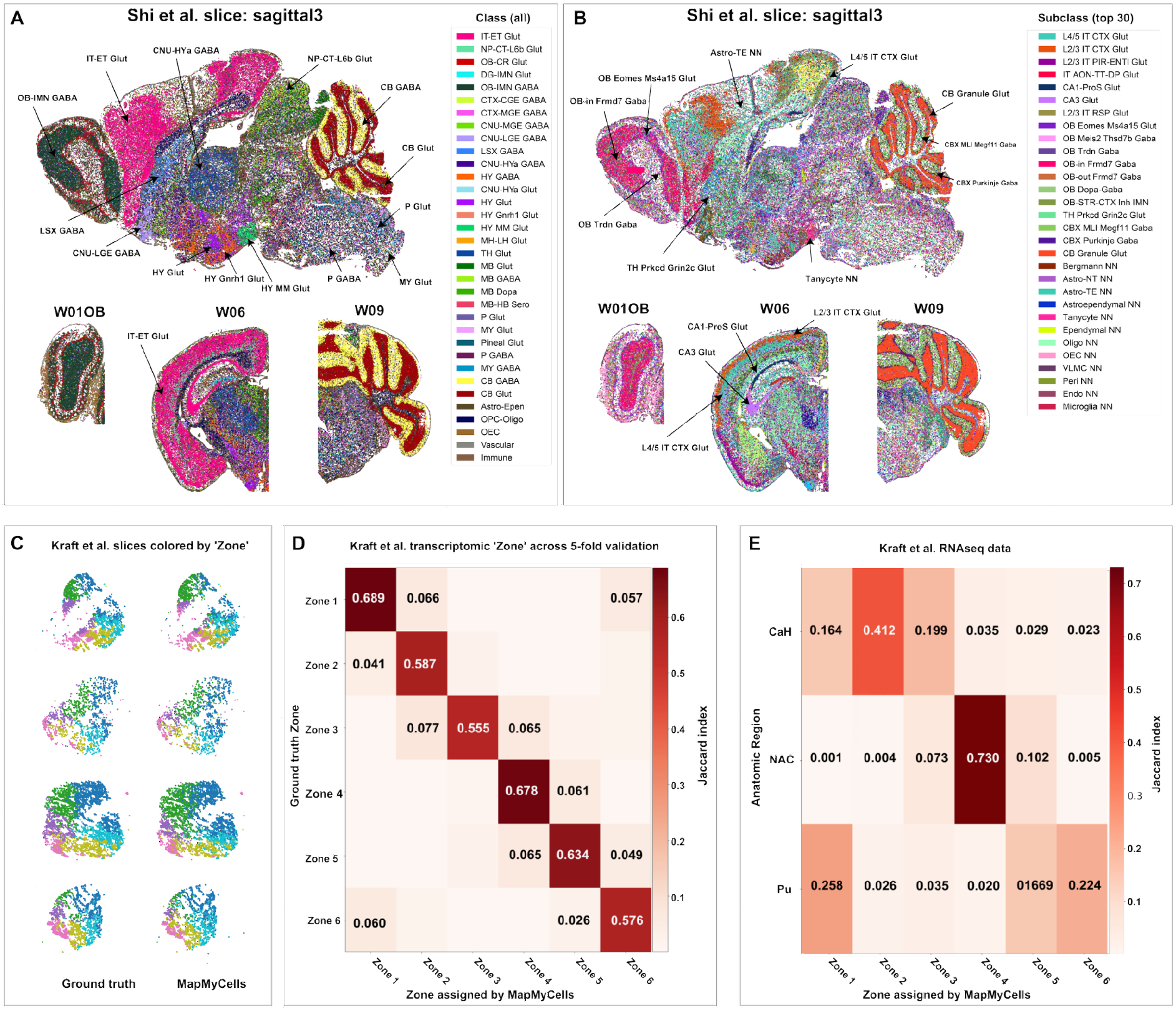
Mapping spatial transcriptomics data. A) and B) representative slices from Shi et al. 2023 spatial transcriptomics data colored according to cell types discovered by MapMyCells. Annotations are class level (A) and subclass level (B). from WMB. C) - E) Results from analysis by Kraft et al. using MapMyCells to assign cells to spatial zones in the human striatum based on their transcriptomic profile. C) Representative slices of spatial data; cells colored according to transcriptomic zone (left), and predicted zones identified by mapping marker genes (right). D) Confusion matrix relating ground truth and mapped “zones” across all iterations of 5-fold validation. E) Confusion matrix relating anatomical dissections from three striatal regions of single nucleus RNAseq data to the transcriptomic “zone” identified by MapMyCells.

#### Assigning cells to spatial zones with MapMyCells

In recent work on molecular zonation of the human striatum across cell types and populations, Kraft et al. ^52^, characterize the cytoarchitectural organization of the human striatum (**Suppl. Fig. 3**). Using an unsupervised method of clustering D1 and D2 matrix medium spiny neurons (MSNs), these cell types were found to exist in six unsupervised clusters, or “zones”, defined by the gene expression data (**Fig. 5C**). The zones have been shown to be conserved across humans in the spatial transcriptomics dataset of 19 donors. These zones are defined by gradients of expression and combinations of gene expression patterns which traverse the striatum in multiple directions, making label transferring these transcriptomically-defined zones challenging. To achieve high-fidelity label-transfer of these zones from one dataset to another, Kraft et al. clustered the zones hierarchically by gene expression similarity, creating a hierarchical taxonomy of zones, as opposed to cell types. Using marker genes derived from this taxonomy, they ran MapMyCells to map the data back into zone space (**Fig. 5C**). Using 5-fold cross validation, they assessed the quality of MapMyCells’ performance based on its ability to preserve the zonal identities of the mapped cells. Aggregated ground truth and all label-transferred zone assignments are shown in **Fig. 5D** and captures by-hand annotation well. After validating the quality of the label transfer on the spatial dataset, Kraft et al. labelled a non-spatial single-nucleus RNA-sequencing dataset of more than 150 donors according to their taxonomy of zones in order to derive a population-level view on gene expression and cell type proportion differences across people. The correspondence of these spatial zones and the reported anatomical region head of caudate nucleus(CaH), nucleus accumbens (NAC), putamen (Pu) of subdissection was calculated. The correspondence between the zone and the canonical anatomical boundaries of the striatum (**Fig. 4E**), indicate quality of the label transfer between the spatial and non-spatial datasets.

## 4. Benchmarking

The expanding repertoire of tools for single-cell omics mapping now enables biologically meaningful integration of new datasets with established cell-type reference frameworks. Numerous methods have been developed for label transfer across single-cell transcriptomic datasets; however, many face scaling challenges when applied to the large cell-by-gene matrices produced by modern high-throughput sequencing platforms^6,53^ . In addition, the absence of definitive biological ground truth complicates evaluation of mapping performance, as existing benchmarks differ widely in task formulation, evaluation metrics, dataset complexity, and output representations.

To perform a limited benchmark for training models we selected a whole mouse cortex and hippocampus taxonomy^54,55^ consisting of 1.17 million cells and clustered to create a taxonomy consisting of 3 classes, 42 subclasses, and 387 clusters. The data was divided into a training set of 820,000 cells and a test set of 350,000 cells. We considered two additional platforms for model comparison, Azimuth^6^, a mutual nearest neighbor method, and CellTypist^9^, a model built on a logistic regression framework. Our attempt to train a model in Azimuth failed during the data integration step due to memory constraints of R when handling very large datasets. With implementation in Python neither CellTypist nor MapMyCells had this difficulty. We trained a CellTypist model in 45 minutes running on a machine with 32 CPUs using 209 GB of memory. The resulting model was configured to label data at the subclass level of the taxonomy. (A model trained to label at the cluster level of the taxonomy required 5.5 hours on 32 CPUs ultimately using 280 GB of memory.) These large requirements are a function of the size of the cell-by-gene matrix.

MapMyCells selects marker genes based on the mean gene expression profile of cell types in the taxonomy. Training MapMyCells required only 4 CPUs and 2.2 GB of memory in 5.5 minutes and was configured to label data at all levels of the taxonomy. Comparing the accuracy of the two methods in **Figure 6A,B**, we find essentially comparable F1 accuracy at the subclass level for the whole cortex dataset (mean MapMyCells 0.96; CellTypist 0.94) and improvement by MapMyCells for mapping at the cluster level (mean MapMyCells 0.75; CellTypist 0.59). This is achieved with significantly lower time and space requirements (**Suppl. Fig. 4A**). A model was also trained at the subclass level using CellTypist for the whole mouse brain taxonomy (WMB)^56^. We find essentially equivalent accuracy of CellTypist and MapMyCells but that performance depends strongly on the 5596 marker gene panel of the unlabeled data. Similar to the analysis underlying **Figure 3B**, we tested CellTypist and MapMyCells’ dependence on gene panel content by constructing test datasets artificially downsampled to small subsets of these 5596 genes, and find that MapMyCells is much more robust against the absence of marker genes than CellTypist (**Fig. 6C**).

**Figure 6.**
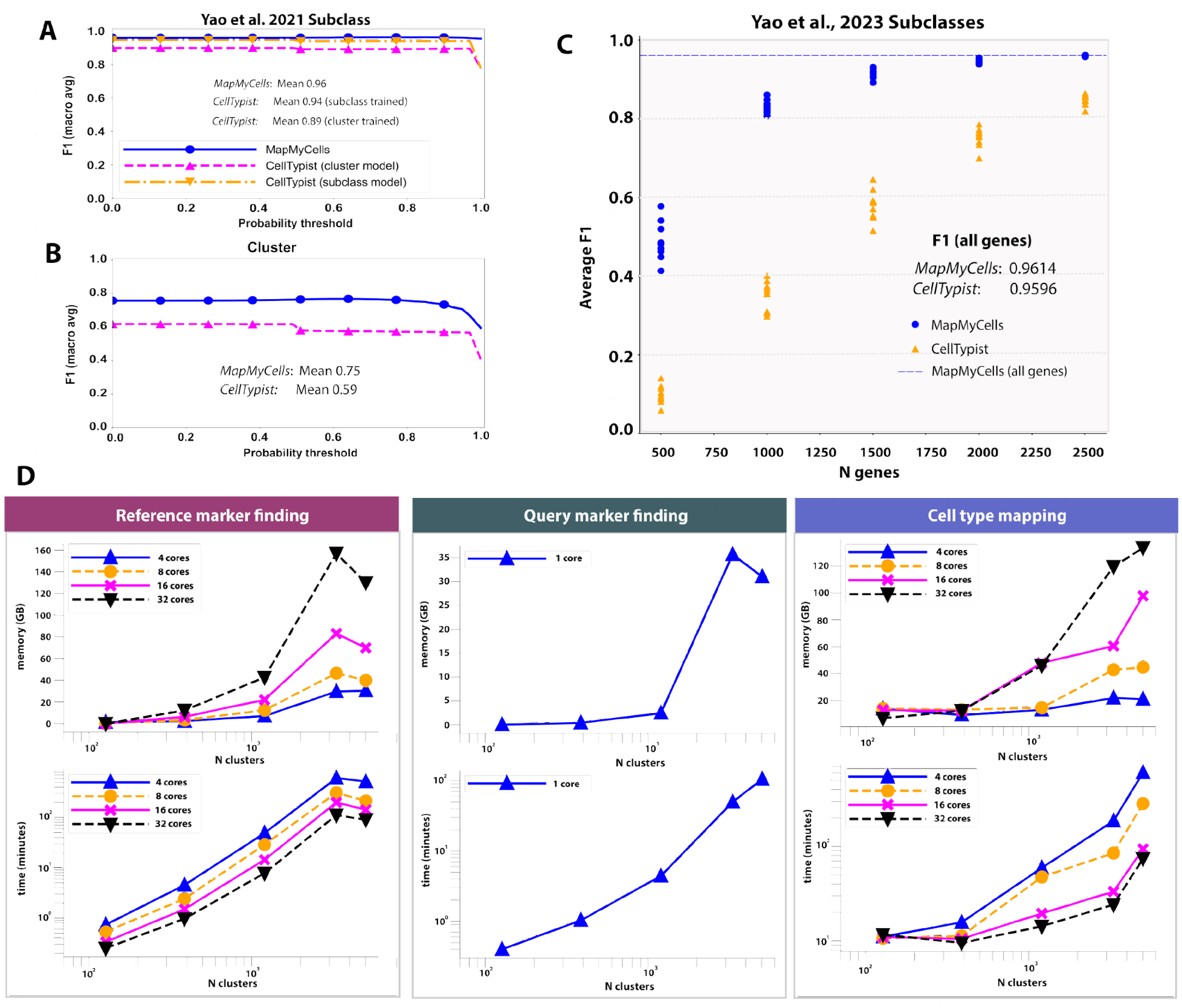
Benchmarking and performance. A) Comparing performance of MapMyCells to CellTypist on the whole mouse cortex taxonomy^54^. F1 accuracy at the subclass level for the whole cortex dataset (mean MapMyCells 0.96; CellTypist 0.94). CellTypist performance slightly degrades when trained at cluster level (mean 0.89). B) Improvement by MapMyCells for mapping at the cluster level (mean MapMyCells 0.75; CellTypist 0.59). This is achieved with significantly lower time and space requirements C) Performance on Whole Mouse Brain (WMB, CCN20230722). MapMyCells and CellTypist achieve comparable overall F1 (mean MapMyCells 0.96; CellTypist 0.95) although CellTypist performance is substantially degraded by limiting the number of training marker genes. D) Resource requirements to map a test set of 350,000 cells to taxonomies of varying complexities with MapMyCells. Curves show requirements by number of cores. The first two columns consider the steps involved in selecting marker genes (see **Methods**). The third column considers the actual process of mapping data to the taxonomy.

### Hardware Requirements

To quantify how MapMyCells’ resource demands grow with the complexity of the taxonomy, we mapped a 350,000 cell by 31,000 gene test dataset onto taxonomies of different sizes on a High Performance Compute cluster managed with the SLURM workload manager^57,58^. By controlling the number of processor cores allocated to each job, we measured the resources in both memory and wall time required to map to a given taxonomy on a given set of hardware. In addition to the taxonomies above, we mapped to much smaller taxonomies (387 clusters) of the mouse cortex and hippocampus^54^ and human primary motor cortex (127 clusters)^29,59^. We also simulated a taxonomy of only 1201 clusters by artificially truncating the WMB taxonomy at the supertype level (the penultimate level of the actual taxonomy).

**Figure 6D** shows the time and memory needed to preprocess and map onto a given taxonomy as a function of the size of the taxonomy (in terms of the number of leaf nodes or “clusters” in the taxonomy) and the number of processor cores devoted to the process. Performance for the three major steps in the MapMyCells algorithm is shown: reference marker selection, query marker selection, and cell type mapping (see algorithm descriptions in **Methods** section). Note that the greedy nature of the query marker selection algorithm makes it difficult to parallelize. For this reason, we only show the performance of query marker finding run on a single core.

In summary, once marker genes have been selected, it is possible to map data onto even the most complex taxonomies on machines the size of a personal laptop given time. Mapping 350,000 cells onto a taxonomy made up of 5,000 clusters requires 20 GB of memory and ten hours of processing on a 4 core machine, 45 GB and 4.7 hours on an 8 core machine. Considering the reference and query marker finding steps, we see that if a user wishes to prepare their own taxonomy for mapping, they can do so in tens of minutes with minimal memory footprint on consumer grade machines for taxonomies with up to ∼ 1,000 leaf nodes. Only the most complex taxonomies (e.g. the Whole Mouse Brain taxonomy with just over 5,000 leaf nodes) require truly high powered computing resources.

### Community Adoption

In October 2023, MapMyCells was made available to the community as a web application^24^. Users can upload either an h5ad^40^ or CSV file of their cell-by-gene data and have their cells mapped to one of the four cell type taxonomies supported by the Allen Institute for Brain Science. Since April 2024, roughly 6400 distinct users have used the MapMyCells web app to map their data. We also track how many cell type annotations have been downloaded from MapMyCells, treating this as a proxy for how many datasets have been mapped using the tool. Since April 2024, 14,600 cell type annotations have been downloaded from the app (**Suppl. Fig. 4B**). MapMyCells has been used to refine the annotation of third-party taxonomies and atlases^60–62^; quantify the link between cell type distribution and disease^63–66^, behavior^67^, development^68,69^ and other epigenomic effects^70^; to assess the targeting specificity of adeno-associated viruses^71,72^; and even to provide training input annotations for machine learning tools^73^.

## Discussion

As the number and complexity of brain cell-type taxonomies continue to grow, there is an increasing need for tools that allow investigators to map arbitrary datasets onto reference taxonomies without dependence on specialized computing hardware. MapMyCells supports scalable mapping of large datasets and complex taxonomies on modest computational workstations and cloud-accessible environments within the Allen Institute Brain Knowledge Platform (BKP). The platform provides accurate mappings across major hierarchical levels of reference taxonomies and can be readily extended to newly developed or evolving taxonomies with minimal preprocessing.

Key determinants of mapping performance include both accuracy and scalability, although accuracy is often difficult to assess rigorously in the absence of independent biological ground truth. Existing tools illustrate these tradeoffs. For example, *Azimuth*^*74*^, a web-based framework built on the Seurat ecosystem, enables mapping of query data to reference taxonomies derived from multiple assay types using a hybrid strategy combining local manifold alignment, k-nearest-neighbor classification, and transfer of labels and embeddings. *CellTypist* employs a multilayer perceptron architecture to map query datasets flexibly to arbitrary taxonomies. While both tools are powerful and accessible, they face limitations in scaling to contemporary dataset sizes. Azimuth reference atlases typically contain on the order of 10^5^ cells and impose limits on the number of query cells that can be mapped per session. CellTypist supports larger references but may require hundreds of gigabytes of memory for model training, restricting practical use to large computing clusters. As reference atlases expand toward millions of cells, such computational requirements are likely to become significant bottlenecks for widespread adoption.

A central design principle of MapMyCells is modularity and interoperability. Marker gene selection is fully decoupled from the mapping procedure, allowing users and consortia to apply independent marker-selection strategies while remaining compatible with shared reference frameworks. The availability of both a web-based graphical interface and an open-source Python library enables broad participation across the neuroscience community and supports standardized label transfer across datasets, species, and experimental modalities.

Algorithmically, MapMyCells operates on mean gene expression profiles of reference cell types. Marker genes are identified through contrasts among leaf-node profiles within the taxonomy, and query cells are mapped by correlating each cell to these profiles within marker-gene space. As a result, computational complexity scales primarily with taxonomy size rather than with the number of cells in either reference or query datasets. This design enables efficient, interpretable, and reproducible mapping of very large single-cell datasets.

We presented several vignettes demonstrating MapMyCells mapping of single-cell RNA-seq, epigenomic ATAC-seq, and spatial transcriptomics datasets. Each modality introduces distinct challenges, including batch effects, platform heterogeneity, biological versus technical variation, mismatches in taxonomy resolution, and mixtures of discrete and continuous cellular states. Cross-modality mapping to epigenomic taxonomies typically requires indirect linkage through gene activity scores, introducing additional uncertainty, while spatial transcriptomic data often contain reduced gene coverage per spot and mixtures of multiple cell types. These challenges suggest that no single mapping method can optimally address all modalities simultaneously.

MapMyCells is therefore designed as a flexible suite of reference taxonomies and mapping methods that prioritize scalability, accessibility, and robust identification of brain cell types across diverse experimental platforms. As the number, diversity, and complexity of cell-type taxonomies continue to expand within the BRAIN Initiative Cell Atlas Network (BICAN), such tools will be increasingly important for enabling community-wide integration of new datasets with shared reference frameworks. By lowering computational barriers and supporting large-scale data integration, MapMyCells aims to facilitate convergence toward consensus cell-type taxonomies for the mammalian brain and to accelerate integration of single-cell omics data with multimodal brain knowledge resources.

## Methods

### MapMyCells Hierarchical Mapping Algorithm

Mapping quality metrics such as the average correlation coefficient for the cells assigned to their final type, and fraction of bootstrap iterations for the assigned cell type are not shown in the pseudocode (**Suppl. Fig. 5**).

#### Algorithm 1 *MapMyCells* (Hierarchical)

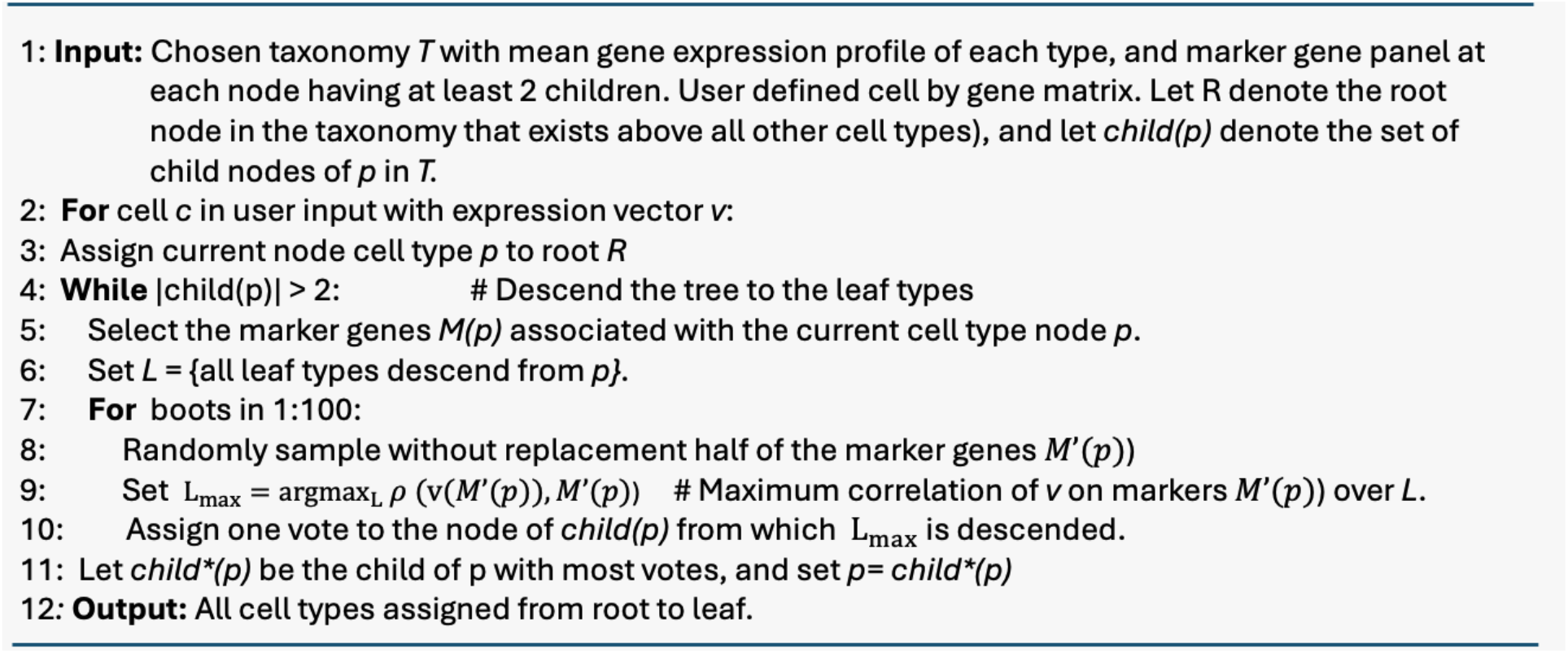

### Quality metrics

Two quality metrics are tracked for each cell type assignment. The “average correlation” is the average of the Pearson’s correlation coefficient (ρ) between the cell and the celltype *averaged over the iterations from line (7) that select that cell type*. The “bootstrapping probability” is the fraction of total iterations that chose the assigned cell type.

### Cell Type Markers

The hierarchical mapping algorithm is agnostic to how marker genes are selected provided input format is respected.^42^ For convenience, the MapMyCells Python library provides functions to select marker genes according to the following algorithm: Marker gene selection is divided into two steps. “Reference marker selection” finds all possible marker genes in the taxonomy. Specifically, it examines every pair of leaf nodes in the taxonomy (e.g. “clusters” in Whole Mouse Brain (WMB) taxonomy) and every gene then denoting each cluster pair, gene) pair with binary “yes” or “no” designation indicating that the gene is or is not a marker when comparing the two clusters. This results in more marker genes than is reasonable to use for hierarchical clustering. “Query marker selection” applies a greedy combinatoric algorithm to downsample this set of marker genes to the much smaller set used by the hierarchical mapping algorithm.

### Reference marker selection

To qualify as a marker gene with respect to a pair of clusters, a gene must pass two filters.The p-value filter applies Student’s t-test to each gene’s expression over the cells assigned to the two leaf nodes to find the p-value representing the likelihood that the gene is distributed differently in the two populations, followed by a Holm-Bonferroni multiple hypothesis correction. Any gene with a corrected P-value < 0.01 passes this filter. The *penetrance P*_*g*_*(t)* of gene *g* in cell type *t* is defined as the fraction of cells in cell type *t* that express gene *g* at greater than 1 count per million. To determine whether *g* is a valid marker in distinguishing two cell types *t*_1_, *t*_2_ we set *q*_*max*_ = *max*(*P*_*g*_(*t*_1_), *P*_*g*_(*t*_2_)), and *q*_*diff*_ = |*P*_*g*_(*t*_1_) − *P*_*g*_(*t*_2_)|/*q*_*max*_. To pass this second filer a cell must satisfy *q*_*max*_ ≥ 0.5 and *q*_*diff*_ ≥ 0.7 and have *log*_2_ fold change greater than unity^75^. This procedure results in the reference marker table RM (as used in query marker selection below).

### Query marker selection

To select the markers distinguishing children of a given taxonomy node the following procedure is used. As with reference marker selection the complexity of the taxonomy is the limiting factor on this algorithm.

#### Algorithm 2

***Query Marker Selection***

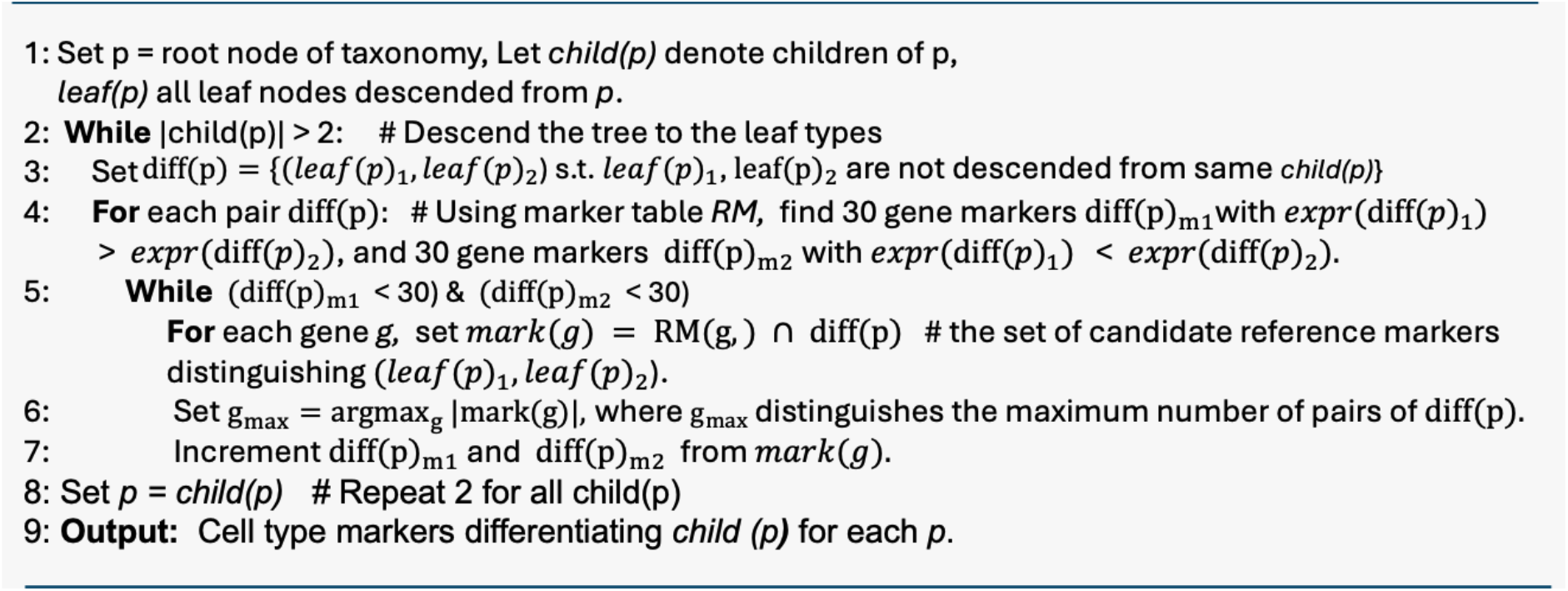

## Acknowledgements

This work was partially supported by NIH grant R01MH123220. The authors would like to thank Mrudhula Balasubramanyan, Eugene Drozd, Arun Dyasani, Jenna Eun, Meenakshi Ponn Shankaran, Ajay Raghunathan, and Vivek Trivedy from Amazon Web Services for designing and building the cloud infrastructure that makes the MapMyCells web app possible.

## Notes

### Competing Interest Statement

The authors have declared no competing interest.

