## Supplemental Figures for "*MapMyCells:* High-performance mapping of unlabeled cell-by-gene data to reference brain taxonomies"

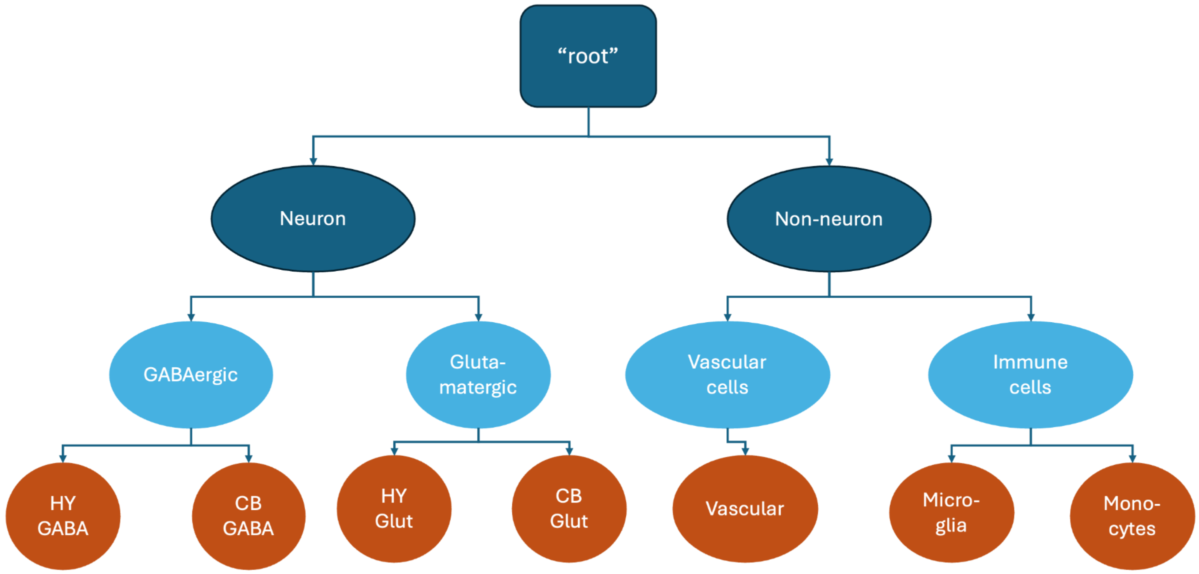


**Supplementary Figure 1.**To map a cell onto this taxonomy, the first step is to assign the cell to the highest level of the taxonomy ("Neuron" or "Non-neuron"). To do this, we take the list of marker genes associated with the "root" node of the taxonomy tree. We randomly subset these markers and use them to correlate the cell's expression against the average expression profiles of the taxonomy's leaf nodes ("HY GABA", "CB GABA", "HY Glut", "CB Glut", "Vascular", "Microglia", and "Monocytes"). Suppose the cell is most correlated with the leaf node "Microglia". This corresponds to one vote for "Non-neuron." We then take a different subset of root-level marker genes and correlate the cell with the leaf nodes again. Suppose that, in this subset of marker genes, the cell is most correlated with the leaf node "CB GABA." This corresponds to a vote for "Neuron". We repeat this process 100 times. Suppose that after those 100 repetitions, we find that there were 65 votes for "Neuron" and 35 votes for "Non-neuron". We determine that the cell is a "Neuron". The next step is to determine whether the cell is "GABAergic" or "Glutamatergic." We take the marker genes associated with "Neuron", subset them, and correlate the cell with the leaf nodes descended from "Neuron" ("HY GABA", "CB GABA", "HY Glut", "CB Glut"). Now, if the cell is most correlated with "HY Glut" that corresponds to a vote for "Glutamatergic," while if the cell is most correlated with "HY GABA", that corresponds to a vote for "GABAergic." Again, we repeat the subset-and-correlate process 100 times. If we then find, for instance, that there were 75 votes for "GABAergic" and 25 votes for "Glutamatergic," we assign the cell to "GABAergic." Finally, to determine whether the cell belongs to "HY GABA" or "CB GABA", we take the marker genes associated with "GABAergic", subset them, and correlate the cell with "HY GABA" and "CB GABA," again repeating the process 100 times, and assigning the cell to the leaf node that received the most votes.

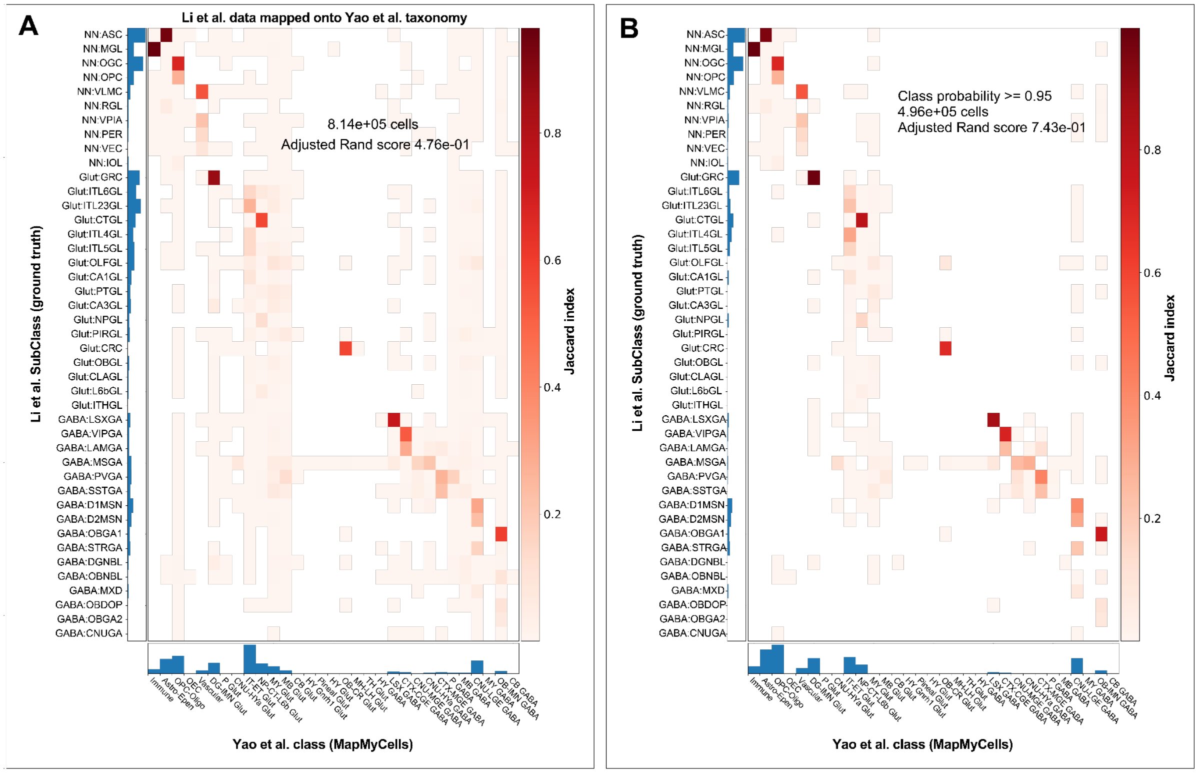


**Supplemental Figure 2.** Applying a quality cut to the mapping result of **Fig 4.** (the green histogram) suppresses the presence of the spurious MY Glut cells resulting in the histogram of **Fig. 4B** and confusion matrices (after quality control cut) relating the cell type WMB classes found by MapMyCells with the subclasses with which Li et al. annotated their data. Salient features here are not only the strong diagonal component (enhanced by the quality cut), but also the biological sensibility of the correspondences. Cells of major classes such as GABAergic, glutamatergic, and non-neuronal are all appropriately assigned.

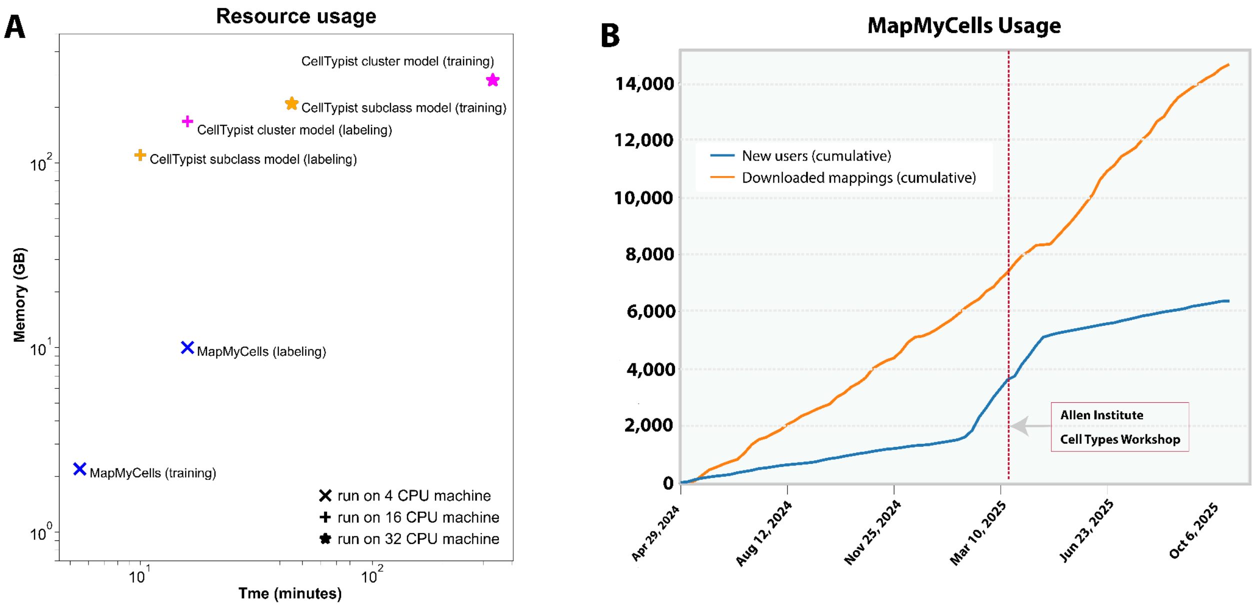


**Supplemental Figure 3.** A) Resource usage comparing MapMyCells and CellTypist over several training and labelling scenarios for 4, 16, and 32 data cores. B) Since April 2024, roughly 6400 distinct users have used the MapMyCells web app to map their data. We also track how many cell type annotations have been downloaded from MapMyCells, treating this as a proxy for how many datasets have been mapped using the tool. Since April 2024, 14,600 cell type annotations have been downloaded from the application.

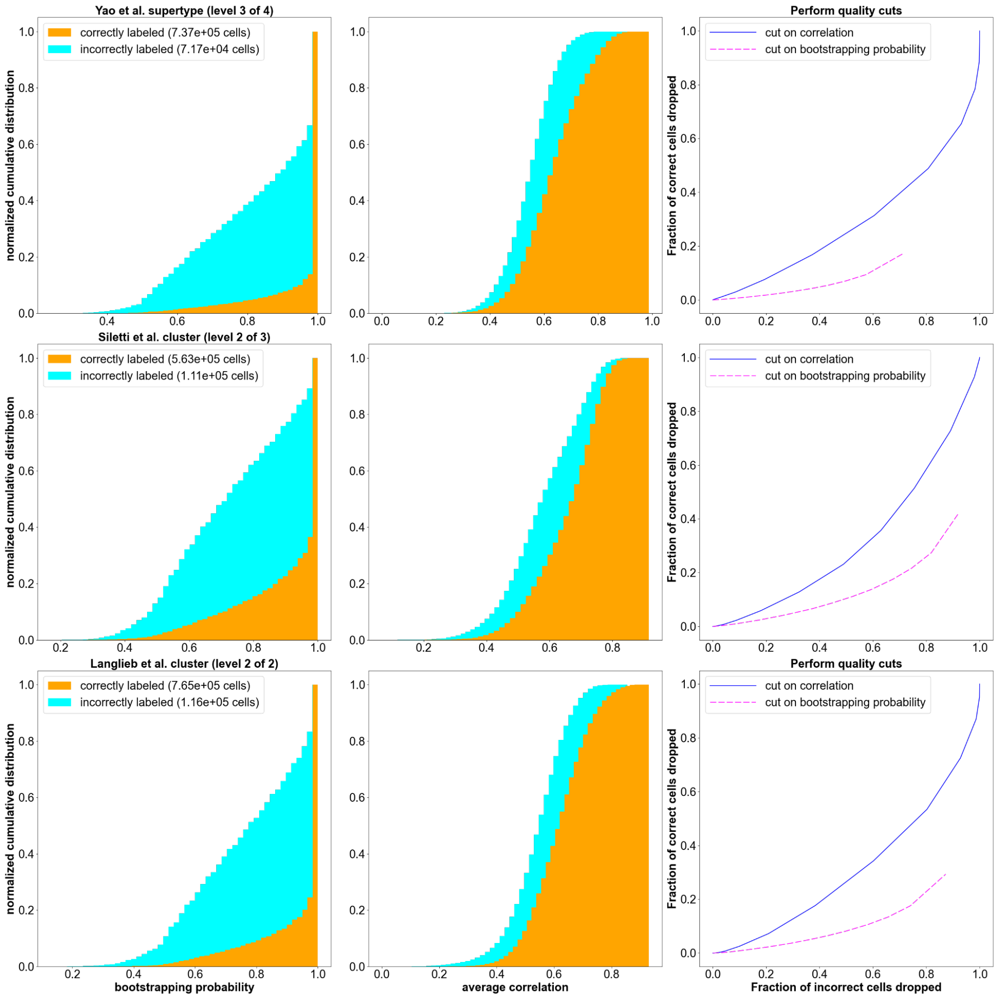


**Supplemental Figure 4.** An examination of the quality metrics returned by the MapMyCells hierarchical mapping algorithm (see **Methods**). For example levels from each of the three taxonomies tested in Figure 3, we show the distribution of "bootstrapping probability" and "average correlation" for both correctly (orange) and incorrectly (cyan) labeled cells. We see that the two distributions differ most significantly in the case of "bootstrapping probability." This has the consequence that quality cuts on "bootstrapping probability" are able to eliminate more incorrectly labeled cells while sacrificing fewer correctly labeled cells than cuts on "average correlation", as demonstrated in the rightmost column.
